# CRISPR-Based Gene Dependency Screens reveal Mechanism of BRAF Inhibitor Resistance in Anaplastic Thyroid Cancer

**DOI:** 10.1101/2025.06.26.661609

**Authors:** Shawn Noronha, Yue Liu, Gaga Geneti, Haojian Li, Xiaolin Wu, David Sun, Vaibhavi Gujar, Takashi Furusawa, Alexei Lobanov, Maggie Cam, Lipika R. Pal, Nishanth Nair, Chi-Ping Day, Eytan Ruppin, Chandrayee Gosh, Jiangnan Hu, Suresh Kumar, Thorkell Andresson, King Chan, Maura O’Neill, Raj Chari, Yves Pommier, Jaydira Del Rivero, Urbain Weyemi, Electron Kebebew, Myriem Boufraqech

## Abstract

Anaplastic thyroid cancer (ATC) is the most aggressive form of thyroid cancer. Despite recent advances in treating BRAFV600E-driven ATC, therapy resistance remains a significant challenge, often resulting in disease progression and death. Leveraging a focused CRISPR/KO screen in parallel with a CRISPR/activation screen, both tailored on response to BRAFV600E inhibitor treatment, we identified TAZ (encoded by the WWTR1 gene) deficiency as synthetically lethal with BRAF inhibitor in ATC. TAZ is overexpressed in ATC compared to well-differentiated thyroid tumors. We demonstrate that TAZ-deficient ATC cells display heightened sensitivity to BRAF inhibitors both *in vitro* and *in vivo*. Using gene essentiality score across a large panel of cancer cell lines, we found that BRAFV600E-driven cancers are highly sensitive to TAZ loss, unlike their counterparts with wild-type BRAF and non-BRAFV600E. Mechanistically, we demonstrate that dabrafenib triggers the Unfolded Protein Response (UPR) under ER stress and suppresses protein synthesis. TAZ loss represses the UPR, reverses the inhibition of protein synthesis, and triggers increased cell death by ferroptosis in dabrafenib-treated ATC. Collectively, our findings unveil TAZ as a new target to overcome resistance to BRAF inhibitors in undifferentiated thyroid cancer.

## Introduction

Anaplastic thyroid cancer (ATC) is a rare yet highly aggressive form of thyroid cancer, comprising less than 2% of all thyroid malignancies. Distinguished by its rapid progression, early metastasis, and poor prognosis, ATC poses significant clinical challenges, with a 1-year survival rate of just 39%. Despite its rarity, ATC accounts for 50% of all thyroid cancer-related deaths [1–3]. The aggressive phenotype and poor prognosis associated with ATC form the basis for its automatic classification as TNM stage IV regardless of tumor burden [4]. Treatment options remain limited and typically involve a combination of surgical intervention, radiation therapy, chemotherapy, and targeted therapies, including BRAF and MEK inhibitors. [5].

The combination of BRAFV600E inhibitor dabrafenib and MEK inhibitor trametinib has demonstrated improved outcome and extended survival in patients with BRAFV600E-mutated anaplastic thyroid carcinoma, offering a significant advancement in the treatment of this aggressive cancer. However, a major challenge persists, as many patients experience severe adverse effects and develop resistance to the therapy, ultimately resulting in disease progression and metastasis. This resistance underscores the urgent need for alternative therapeutic strategies to provide sustained responses in combination with BRAF inhibitors and further improve patient outcome.

Here, we developed focused CRISPR knockout and activation libraries comprising ∼2,000 genes differentially regulated upon BRAFV600E inhibitor dabrafenib treatment. These libraries were harnessed to screen for mechanisms of resistance to targeted therapy in ATC cells. Our screening identified *WWTR1* (WW domain containing transcription regulator 1), the gene encoding transcriptional coactivator with PDZ-binding motif (TAZ), as a critical driver of resistance to dabrafenib in ATC cells.

TAZ is a key effector of the Hippo signaling pathway. Lacking a DNA-binding domain, TAZ functions as a co-regulator by interacting with DNA-binding proteins to modulate gene transcription. Transcriptional enhanced associate domain (TEAD) transcription factors are the primary partners of TAZ and its paralog Yes-associated protein (YAP) in driving transcriptional activity. In the Hippo pathway, the MST1/2 kinases phosphorylate the LATS1/2 kinases, which in turn phosphorylate YAP and TAZ, leading to their cytoplasmic retention and subsequent degradation. When the Hippo pathway is inactive, dephosphorylated YAP and TAZ can translocate to the nucleus and drive transcription via binding partners like TEADs [6–8]. Recent studies uncovered the distinct activities and non-redundant functions of TAZ and YAP in carcinogenesis and response to therapy [9]

Endoplasmic reticulum (ER) stress, caused by the accumulation of improperly folded proteins, triggers an adaptive response known as the unfolded protein response (UPR). The UPR aims to alleviate misfolded protein accumulation by attenuating protein translation and restoring protein homeostasis. Increased expression of protein sensors involved in the UPR has been observed in various human tumors. UPR activation occurs when the chaperone heat shock protein family A (HSPA5) dissociates from ER stress sensors, including protein kinase RNA-like ER kinase (PERK), inositol-requiring enzyme 1 alpha (IRE1α), and activating transcription factor 6 alpha (ATF6α), leading to their autophosphorylation and activation of downstream signaling pathways [10, 11]. Recent studies have linked enhanced tumor therapy resistance to ER stress and the activation of the UPR, an adaptive response aimed at restoring cellular homeostasis and preventing cancer cell death [12–14].

In this study, we demonstrated that TAZ depletion sensitizes ATC cells to the BRAF inhibitor dabrafenib, reduces tumor growth, and prolongs survival *in vivo*. We further showed that dabrafenib activates the UPR, increases activating transcription factor 4 (ATF4) expression, and upregulates its downstream target phosphoglycerate dehydrogenase (PHGDH), which is essential for de novo serine synthesis. Notably, TAZ inhibition during dabrafenib treatment reverses the adaptive UPR, partially restores abnormal protein synthesis, and enhances ferroptosis.

Our findings unveil TAZ as an actionable therapeutic target to overcome resistance to BRAF inhibitors in undifferentiated thyroid cancer.

## Results

### Focused CRISPR/KO and CRISPR/Act screens identify *WWTR1* (TAZ) as a critical driver of resistance to dabrafenib in anaplastic thyroid cancer

To comprehensively investigate the genetic determinants of resistance to BRAFV600E inhibitors, we first compiled a list of genes in BRAFV600E-mutated cell lines that exhibit a negative CHRONOS gene effect score in the Cancer Dependency Map (DepMap) compared to Non-BRAFV600E and Wild type-BRAF cell lines. This list was then merged with a set of differentially expressed genes identified in dabrafenib-treated cells through RNA sequencing of 8505c cells treated with dabrafenib for 14 days. After integrating these gene lists and excluding noncoding RNAs (ncRNAs) and pseudogenes, we constructed a focused CRISPR gene library comprising 2,050 genes for use in knockout (KO) and activation (act) screens. We then generated a CRISPR knockout (KO) and CRISPR activation libraries, each comprising six guides per gene for 2050 genes. Each library was transduced into Cas9 and dCas9/MPH positive 8505c cells respectively. After 7 days of puromycin selection, cells were treated with a concentration of dabrafenib corresponding to the ≈IC20 (**Supplementary Fig 1A).** At 7 and 14 day treatments, surviving cells were harvested and gRNA barcodes identified by next-generation sequencing. Using four (4) replicates per treatment group, the relative depletion or enrichment of the guides in drug treated groups was compared to the control cells using *MAGeCKFlute* differential Beta scores (**Fig 1A**).

**Figure 1.**
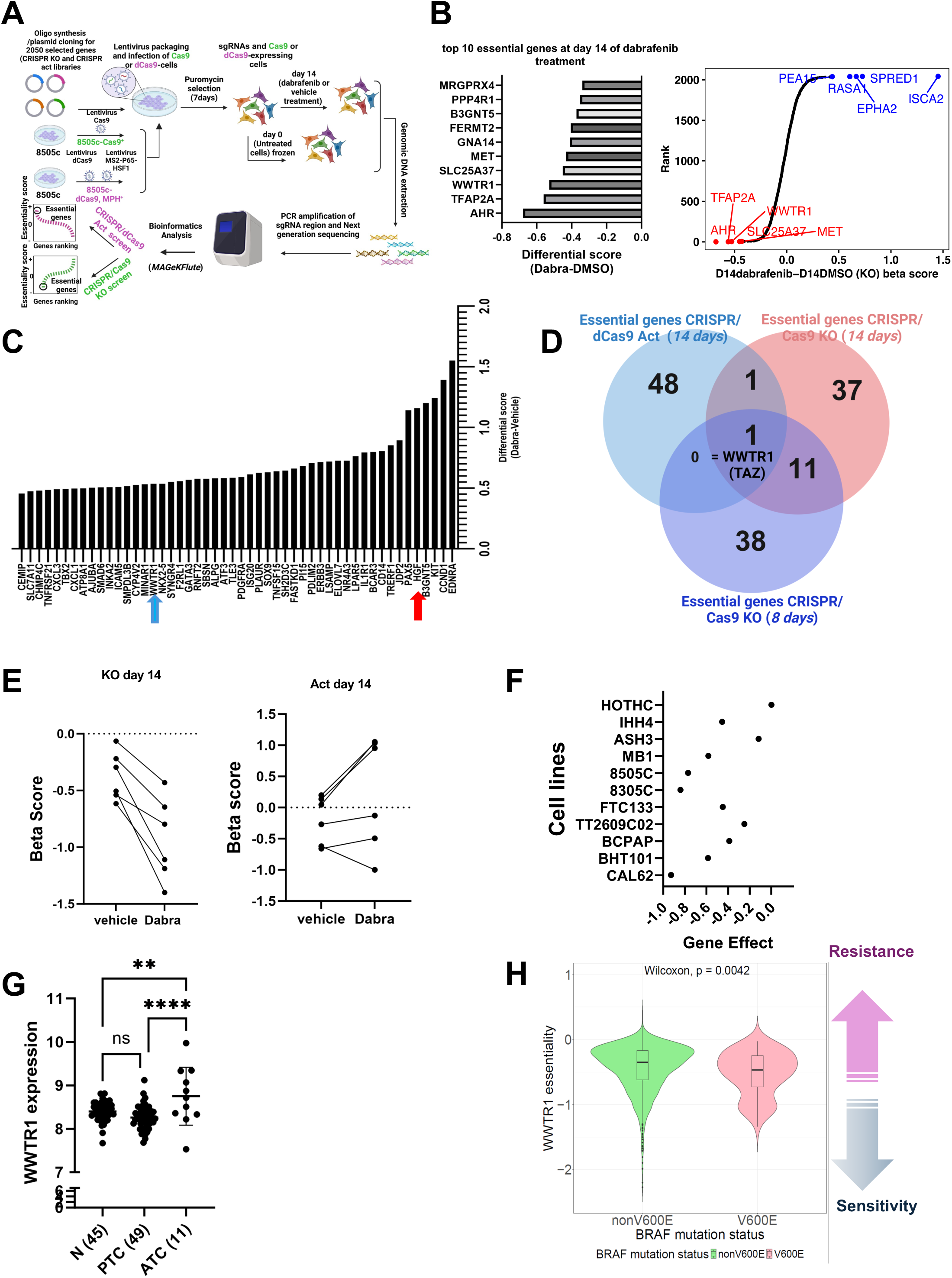
Focused CRISPR/KO and CRISPR/Act screens identify WWTR1 (TAZ) as a critical gene for resistance to dabrafenib in anaplastic thyroid cancer. **A.** Human focused CRISPR/Cas9 KO library and CRISPR/dcas9 activation library containing number of guides were packed into lentiviral particles and transduced into Cas9+ or dCas9+ 8505c cells. The transduced cells were selected with puromycin. Newly generated mutant cell lines were cultured for 14 days with vehicle or dabrafenib (0.4 µM) for screening. Genomic DNA was extracted from the treated cells with vehicle or dabrafenib, sgRNA fragments were amplified by RT-PCR followed by next-generation sequencing. Genomic DNA from non-treated 8505c-transduced cell line was used as control. Sequencing data were analyzed using MAGeCKFlute. Genes were ranked by essentiality. **B**. Top 10 hits from the CRISPR/Cas9 KO at 14 days of dabrafenib treatment based on differential beta score**. C.** Top 50 “essential genes” from CRISPR/dCas9 activation screen. Red arrow pointing to HGF and blue arrow pointing to WWTR1. **D**. Venn Diagram showing the common essential genes from the CRISPR/Cas9 KO and activation screens’ top 50 hits. **E.** sgRNA targeting WWTR1 (TAZ) were depleted in dabrafenib treated cells and sgRNA activating WWTR1 were enriched in dabrafenib treated cells, suggesting that WWTR1 is an essential gene for survival during treatment. **F**. Cancer Dependency Map showing gene effect of WWTR1 using CHRONOS score in thyroid cancer cell lines**. G**. Gene expression Omnibus dataset (GSE33630) showed higher expression of WWTR1 in ATC compared to normal thyroid or PTC. **H**. WWTR1 (TAZ) essentiality in cancer cell lines based on BRAF mutation status using DepMap dataset. Essentiality score is defined by the effect of WWTR1 loss on cell viability. BRAFV600E-driven cancer cells are more sensitive to WWTR1 KO (TAZ KO) than non-BRAFV600E cells. *p<0.05, **p<0.01, ***p<0.001, ****p<0.0001

Among the top fifty essential hits from the CRISPR screens, only WWTR1 guides were lost at 7 days and 14 days in dabrafenib-treated cells transduced with the CRISPR KO library and enriched in dabrafenib-treated cells transduced with the CRISPR activation library (**Fig 1B, C, and D, Supplementary Fig 1B)**). All 6 gRNAs targeting WWTR1 were negatively enriched in dabrafenib-treated cells transduced with the KO library (**Fig 1E, left panel, Supplementary Fig 1C**), and positively enriched in dabrafenib-treated cells transduced with the CRISPR activation when comparing cells treated with the drug to their vehicle counterparts (**Fig 1E right panel**).

MET, one of the top candidates from the CRISPR KO screen (**Fig. 1B**), and its ligand HGF (red arrow), a top candidate from the CRISPR/dCas9 activation screen **(Fig. 1C),** have been shown to play a role in resistance to BRAF inhibition in a murine ATC model [20]. These findings further validate our screening results. However, while our data revealed Aryl Hydrocarbon Receptor (Ahr) as potentially the most required gene for survival upon dabrafenib treatment in the CRISPR/Cas9 KO screen, AhR depletion led to increased cell proliferation in the presence of dabrafenib. This observation was further supported by the *DepMap* CRISPR/Cas9 KO data across a large panel of thyroid cancer cell lines (**Supplementary Fig. 1D, 1E**).

### Inhibition of WWTR1 (TAZ) sensitize ATC cells to dabrafenib

To delineate the role of WWTR1 gene, which encodes for TAZ protein, in the sensitivity to dabrafenib; we used publicly available databases, along with shRNA-depletion and KO cells. Examination of genome-wide CRISPR/Cas9 KO screen results for thyroid cancer cells from *DepMap* showed reduced cell growth in TAZ KO cells, revealing that TAZ is an essential gene for survival in ATC cells **(Fig 1F).** The analysis of WWTR1 expression in different histological types of thyroid cancer revealed a higher expression in human ATC tumors compared to differentiated thyroid cancer (PTC) and normal tissue **(Fig 1G).** Similarly, the essentiality score, which is the normalized growth reduction resulting from gene inactivation, across all cancer cell lines showed a negative essentiality score for WWTR1 in BRAFV600E mutant cancer cells compared to the non BRAFV600E cells (one-sided Wilcoxon rank-sum test, p = 0.0042; n = 68 BRAF V600E cell lines, n = 67 other BRAF mutant cell lines, n = 941 BRAF wild type cell lines; Lee et al., 2021). This data suggests that there is a higher sensitivity to the loss of WWTR1 in BRAFV600E-driven cancer cell lines compared to the non-BRAFV00E cancer cell lines (**Fig 1H)**. Altogether, these data suggest that WWTR1 is required for survival of cells with oncogenic BRAF.

To further validate the results from the gene essentiality and CRISPR screens, we generated TAZ depleted-8505c cells using shRNA and independent sgRNAs targeting TAZ. TAZ-depleted cells were treated with increasing concentrations of dabrafenib, followed by measurement of viability and cell growth. Loss of TAZ significantly sensitizes ATC cells to dabrafenib as measured in cell growth and colony formation assays **(Fig 2A-C top and bottom panel,**). Specifically, measurement of cell death in clonogenic assays revealed over 60 to 70% of cell death upon dabrafenib following TAZ depletion, while only ∼28-30 % of cell death was detected with dabrafenib treatment alone. Of note, the loss of TAZ alone only exhibited 30% effect on cell viability in the absence of BRAF inhibitor. Collectively, these data suggest that TAZ is a key driver of resistance to BRAF inhibitor in ATC cells. Stably transduced cells were confirmed by immunoblotting and RT-PCR (**Fig 2D**).

**Figure 2.**
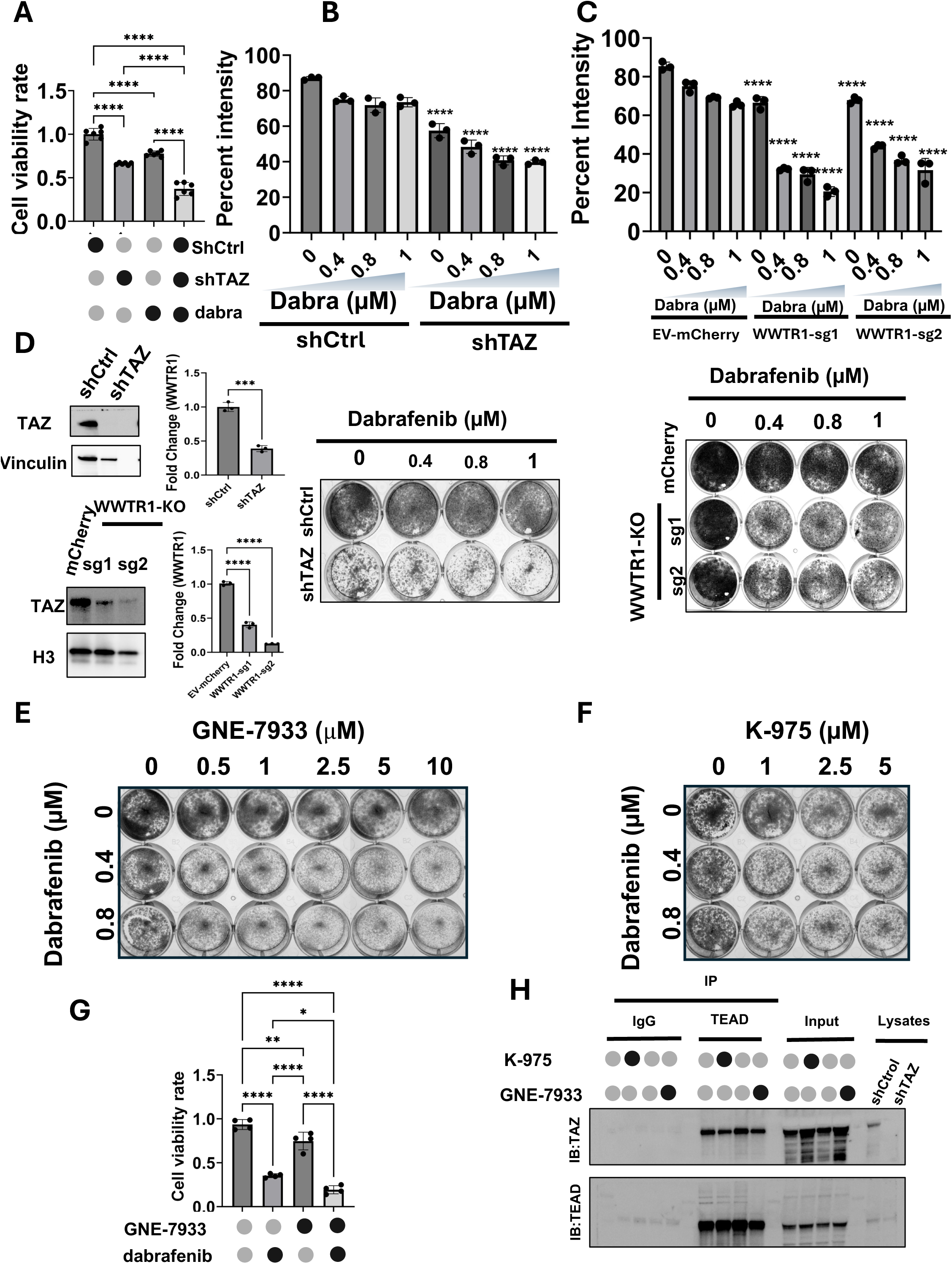
Inhibition of TAZ (encoded by WWTR1) sensitize BRAF^V600E^-driven ATC cells to dabrafenib. **A.** Depletion of TAZ using shRNA showed a mild effect on cell viability, However, it significantly reduced cell growth in the presence of dabrafenib **B**. Top and bottom panel, Colony formation assay showing that TAZ depletion sensitizes ATC cells to dabrafenib. **C**. TAZ-knock out markedly reduces colonies formation upon dabrafenib treatment**. D.** TAZ expression by western blot and RT PCR in shTAZ cells, and TAZ KO cells using CRISPR/Cas9 system and sgWWTR1-sg1 and sgWWTR1-sg2. **E, F**. TAZ-TEAD inhibitors, GNE-7933 and K-975 sensitized 8505c to dabrafenib. **G**. ATC patient-derived spheroids showed a significant reduction of cell viability in cell spheroids treated with the combination of GNE-7933 and dabrafenib compared to single agents. **H**. Immunoprecipitation assay showed a reduction of the binding of TAZ to TEAD with K-975 and GNE-7933. *p<0.05, **p<0.01, ***p<0.001, ****p<0.0001.

The main oncogenic function of TAZ has been attributed to its binding to the transcription factor TEAD. We then asked whether the inhibition of TAZ-TEAD interaction using two pharmacological inhibitors, K-975 and GNE-7933, mimics the synthetic lethality of TAZ depletion with dabrafenib. Increasing concentration of K-975 and GNE-7933 successfully sensitized ATC cells to dabrafenib (**Fig 2E, F, Supplementary Fig 2A**). A similar anti-proliferative effect was observed in spheroids derived from a patient with BRAF^V600E^-driven anaplastic thyroid cancer (ATC01) (**Fig 2G)**. These observations are consistent with the ability of these drugs to disrupt TAZ and TEAD interaction as shown in **Fig 2H**. As the interaction between TAZ and TEAD was partially disrupted in the presence of either K-975 or GNE-7933, this effect was more pronounced when these inhibitors were combined with dabrafenib (**Supplementary Fig 2B)**.

### TAZ depletion overcomes resistance of ATC tumors to dabrafenib in mice and prolongs survival

We next validated the synthetic lethality of TAZ depletion and dabrafenib treatment using a xenograft model. TAZ-depleted (shTAZ) or control 8505c tumor cells (shCtrl) were injected into the flank of immunocompromised animals and treated daily with 15mg/kg of dabrafenib or vehicle control. Dabrafenib-treatment led to a 30% reduction in tumor growth *in vivo*, while TAZ depletion did not show a significant change. In contrast, TAZ depletion upon dabrafenib treatment exhibited a 53% diminution in tumor size compared to the control mice (**Fig 3A)**. These observations were also substantiated by a significant reduction in tumor weights ex vivo **(Fig 3B)**. Noticeably, dabrafenib treatment did not yield adverse effects, nor changes in body weight throughout the treatment, implying that the drug was relatively well tolerated (**Fig 3C**). Furthermore, the animal group implanted with TAZ-depleted cells treated with vehicle, and the group injected with control cells treated with dabrafenib showed a modest improvement in median survival (14 days for dabrafenib treated group compared to 7 days for the control group). Most remarkably, TAZ depletion led to more than 3-fold increase in survival when mice were treated with dabrafenib (median survival of 25.5 days), with about 30% of the animals surviving past the study endpoint (**Fig 3E, 3F**).

**Figure 3.**
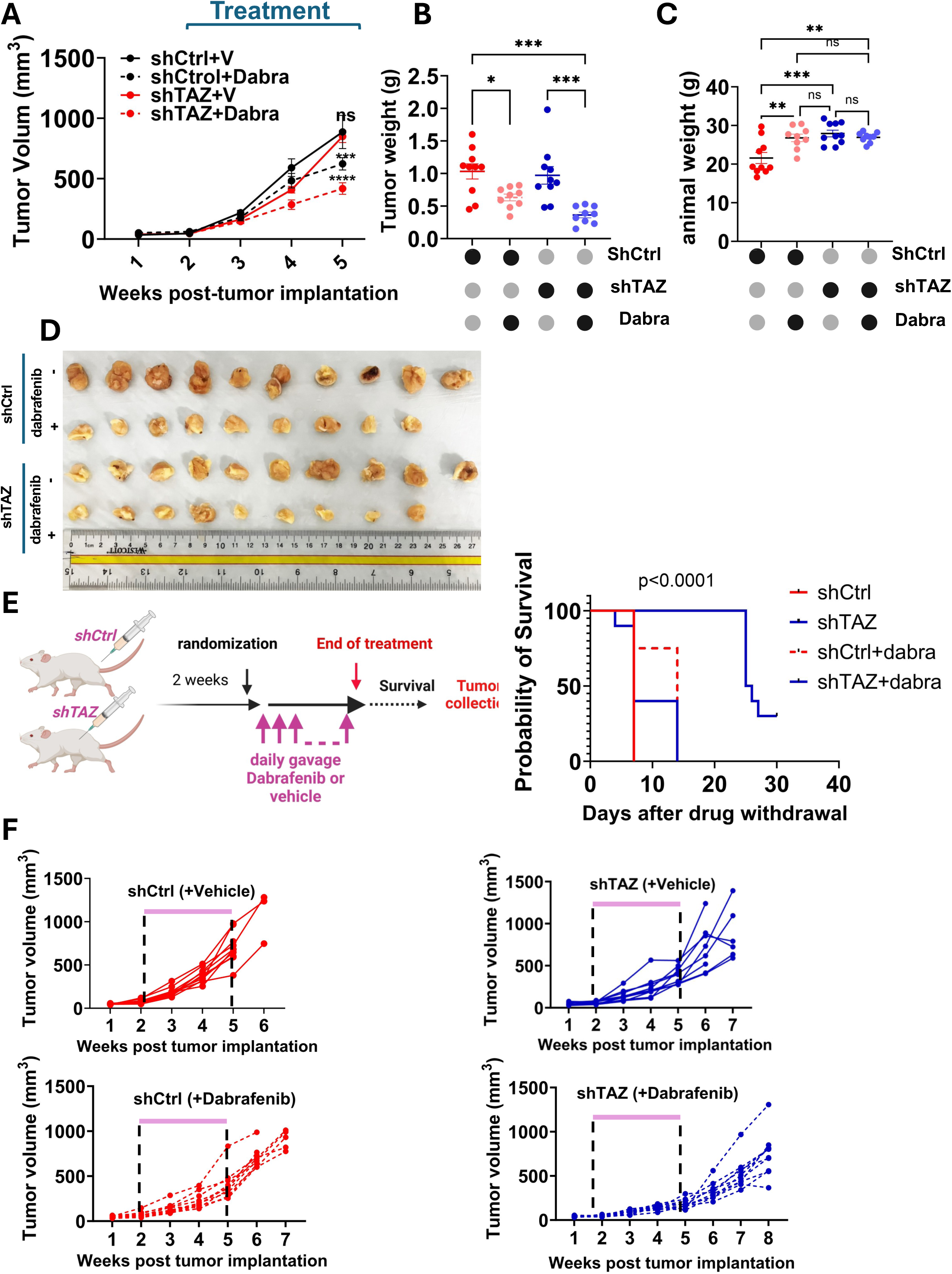
TAZ depletion sensitizes tumor xenografts to dabrafenib and prolong survival. **A.** Tumor growth curves of tumor xenografts in immunocompromised mice. 8505c-shCtrl or 8505c-shTAZ cells were implanted subcutaneously into the flanks. Treatment with vehicle or dabrafenib was initiated 2 weeks after tumor implantation. Dabrafenib was administered daily by gavage. Tumor diameters were measured weekly and used to calculate tumor volume. Data show mean±SEM (*n* = 8 or 10). **B**. Tumor weight was recorded ex vivo. **C**. Animal weight recorded at the end of the experiment showing no weight loss due to dabrafenib treatment. **D**. Ex vivo images of the tumors harvested at the end point of the experiment **E.** Kaplan Meier survival curve after dabrafenib withdrawal. **F**. Growth curve over time of each tumor implanted for up to 3 weeks post drug withdrawal. *p<0.05, **p<0.01, ***p<0.001, ****p<0.0001

### TAZ inhibition antagonizes dabrafenib-induced ER stress response

To investigate the molecular mechanisms underlying TAZ-driven resistance to dabrafenib, we analyzed global transcriptomic and proteomic changes in dabrafenib-treated and TAZ-depleted cells. Gene set enrichment analysis revealed a prominent upregulation of the unfolded protein response (UPR) genes in cells treated with dabrafenib (**Fig 4A**). Dabrafenib treatment reactivates the expression of the genes coding for key proteins of the UPR (**Fig 4B**), genes which were reduced with TAZ depletion (**Fig 4C**). Additionally, proteomic analysis with a false discovery rate (FDR) cutoff of 0.05 revealed an activation of a set of proteins involved in the UPR and in amino acid synthesis and transport, such as SLC7A5, ASS1, SLC1A5 and PHGDH, in dabrafenib-treated cells compared to controls (**Fig 4D**), supporting our transcriptomic analysis showing activation of the UPR in response to dabrafenib.

**Figure 4.**
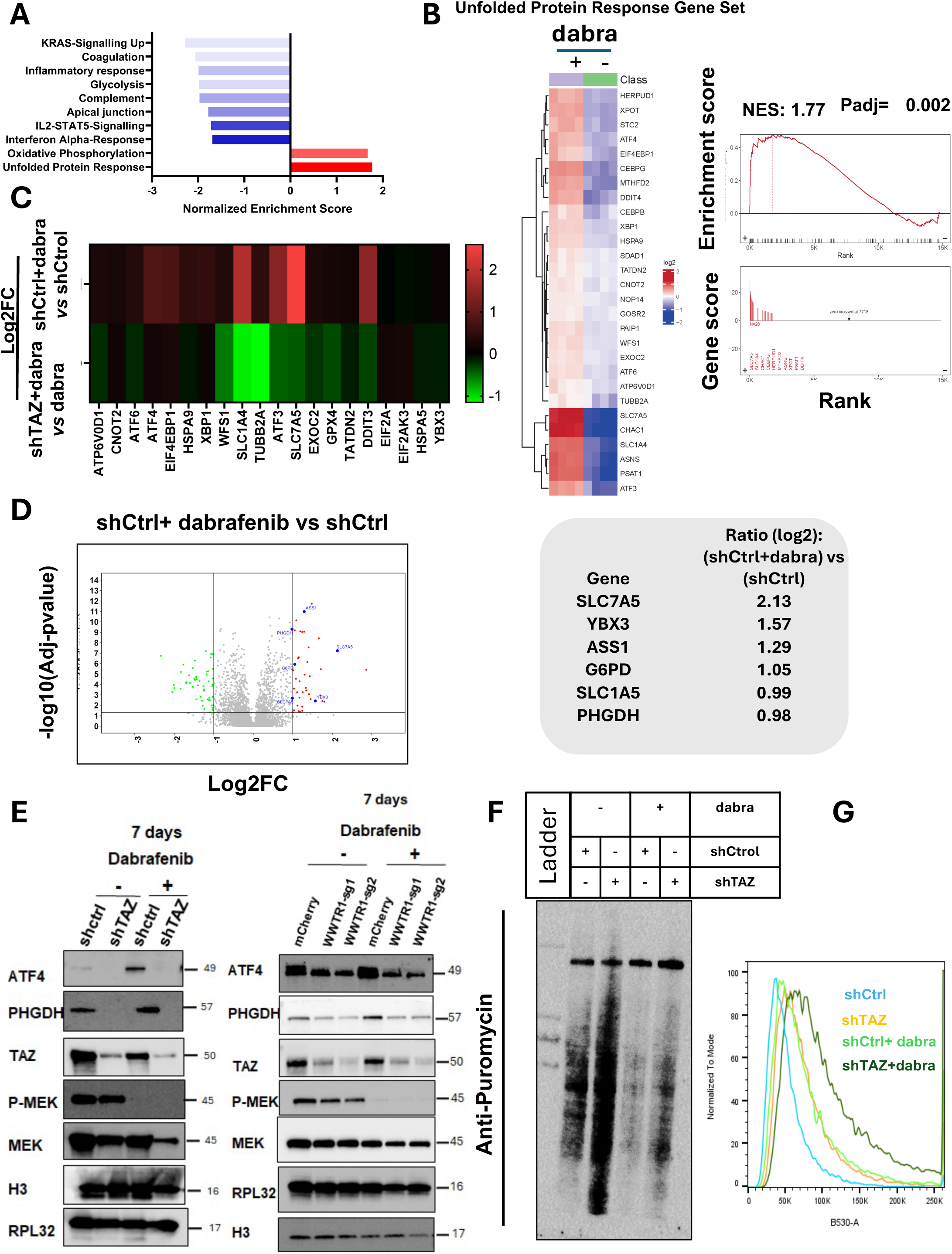
Loss of TAZ reverses the dabrafenib-induced UPR in BRAFV600E-driven ATC cells. **A.** Gene Set Enrichment Analysis (GSEA) for top 10 pathways showing that the unfolded protein response (UPR) is the most enriched pathway in response to dabrafenib treatment. **B.** Heat map showing the expression profile of the UPR genes in dabrafenib and vehicle treated control cells. **C**. Heatmap of log2fold-change value of differentially expressed genes of dabrafenib *vs* vehicle and dabrafenib treated shTAZ cells *vs* dabrafenib treated shCtrl cells. **D**. **Left panel** Proteomics profiling of dabrafenib treated cells. Difference is plotted in x-axis and −log10(adj P-value) on the y-axis. **Right panel**. Downstream effectors of UPR in dabrafenib treated cells. **E.** Cells depleted for TAZ (shTAZ and TAZ KO) were treated with dabrafenib for the indicated times. ATF4, PHGDH and p-MEK were analyzed. TAZ depletion was confirmed in shTAZ and TAZ KO cells. H3 and RPL32 were used as loading controls **F**. Analysis of newly synthesized proteins using puromycin incorporation for 10min at 7 days of dabrafenib treatment in the presence or absence of TAZ**. G.** Lipid ROS level was analyzed using BODIPY in shCtrl, shTAZ, shCtrl+dabrafenib and shTAZ+dabrafenib cells. RSL3 was used as a positive control. Concentration of dabrafenib used is 0.8 µM.

Activation of the UPR triggers two distinct cellular events to mitigate the ER stress: first, a reduction of newly synthesized proteins and enhanced degradation of misfolded proteins, followed by a transcriptional increase of set of genes involved in amino acid synthesis, transport, and regulation of proteostasis [11, 21].

Based on these transcriptomic and proteomic observations, we hypothesized that dabrafenib-induced ER stress response requires TAZ, and that TAZ deletion leads to impaired protein synthesis and cell death. To validate this hypothesis, TAZ-depleted cells and TAZ knockout cells were treated with dabrafenib and analyzed for UPR protein ATF4 and its downstream effectors. The UPR induces metabolic rewiring via serine biosynthesis [22]. As the genes of the serine biosynthesis pathway (SSP) are transcriptionally regulated by ATF4.

Dabrafenib treatment significantly upregulated ATF4 and PHGDH, in a dose-dependent manner (**Supplementary Fig 3A**). Phosphoglycerate dehydrogenase (PHGDH), the rate limiting enzyme in serine biosynthesis, has been classified as a potential oncogene [23, 24]. Moreover, serine-derived glycine is a component of GSH, thus SSP can regulate cell survival by participating in redox reactions and redox homeostasis. [25] [26] [27]

Depletion of TAZ markedly suppressed the expression of both proteins as early as 72 hours, and this effect persisted even in the presence of dabrafenib **(Supplementary Fig 3B,-C Fig 4E)**. TAZ-TEAD inhibition had a similar effect on PHGDH and ATF4 in the presence of dabrafenib **(Supplementary Fig 3D)**. We confirmed the role of TAZ in UPR using the ER stress inducer, Thapsigargin (TG). Thapsigargin treatment increased both TAZ and ATF4 expression **(Supplementary Fig 3E)**. Analysis of the two other main regulators of the SSP, PSAT1 and PSPH, by RT-PCR at 7 days of treatment showed a similar profile as PHGDH (Supplementary Fig 3F). ATF4 activates the transcription of the SSP genes in response to stress induced-amino acid depletion [28]. To test further whether the SSP inhibition observed can be mediated by the loss of ATF4, cells were inactivated for ATF4 using small interference RNA. ATF4 knockdown reduced *PHGDH*, albeit with no effect on *WWTR1* (TAZ transcript) (**Supplementary Fig 3G**).

Under ER stress conditions, the protein chaperone HSPA5 dissociates from ER stress sensors, thus leading to their autophosphorylations and subsequent activation of downstream pathways [29, 30]. PERK, a key ER sensor, oligomerizes and transphosphorylates to inhibit protein translation, thus reducing the newly synthesized protein entering the ER. This allows for a selective activation of ATF4 transcription. Gene expression analysis in a cohort of patients with thyroid cancer showed a significant positive correlation between WWTR1 and ATF4 (Spearman correlation r=0.33, p<0.001) (**Supplementary Fig 3H)**. Taken together, our findings strongly suggest that the UPR protects ATC cells from chronic ER stress induced by BRAFV600E inhibition, thus leading to drug resistance. TAZ loss impairs this protective mechanism, creating a genetic vulnerability in ATC cells.

To determine the impact of pharmacological inhibition of BRAFV600E and TAZ depletion on protein translation during the ER stress and ATF4, a puromycin incorporation assay was used at 7 days of treatment with dabrafenib. Loss of ATF4 in TAZ-depleted cells was associated with a pronounced increase in protein synthesis (**Fig 4F)**. Notably, the elevated rate of ATF4 upon dabrafenib treatment is associated with inhibition of protein synthesis, thus suggesting that the UPR induced by dabrafenib is a mechanism of resistance to BRAFV600E inhibition (**Fig 4F**). More importantly, TAZ depletion in dabrafenib-treated cells partially restored protein synthesis (**Fig 4F**). These data suggest that TAZ depletion and the subsequent downregulation of ATF4 in dabrafenib-treated cells partially restores protein synthesis under ER stress conditions. Thus, in response to dabrafenib, we propose that TAZ hijacks the UPR to maintain cellular homeostasis under stressful conditions. Conversely, TAZ depletion disrupts the UPR, resulting in chronic stress and ultimately leads to cell death.

### TAZ depletion promotes ferroptosis in dabrafenib-treated ATC cells

We next examined the mechanisms underlying cell death in TAZ-depleted cells upon the pharmacological inhibition of oncogenic BRAF. The ER stress-driven UPR, and ATF4 activation enable the HSPA5-GPX4 complex to prevent ferroptosis [31]. Similarly, PERK-ATF4-SLC7A11 axis can modulate cancer cell death by ferroptosis [32]. Cell death analysis showed that silencing of TAZ upon BRAFV600E inhibition significantly reduces caspase3/7 activity (**Supplementary Fig 4A).** Analysis of ferroptosis by quantification of membranes lipid peroxidation using BODIPY-C11 probe showed a two-folds increase in ferroptosis in TAZ-depleted cells treated with dabrafenib (**Fig 4G, Supplementary Fig 4B**). These observations are corroborated by the evidence of oxidative stress in the same cells, as revealed by *CellRox* Green fluorescence levels **(Supplementary Fig 4C)**. These findings are consistent with previously published data [32, 33], indicating that the UPR triggers ferroptosis when ER stress is maintained due to lack of ATF4 and partial recovery of protein synthesis despite a persistent protein misfolding.

### PHGDH or PERK inhibition mimic TAZ depletion in sensitizing ATC cells to dabrafenib

Because PHGDH, the rate limiting enzyme in the *de novo* SSP, is reduced in TAZ-depleted cells treated with dabrafenib. We examined whether pharmacological inhibition of PHGDH can mimic the synthetic lethality of TAZ depletion with dabrafenib. Colony formation assay showed a synergistic effect of dabrafenib and the PHGDH inhibitor CBR-5884 on ATC cells growth compared to the effects achieved by single agents at doses demonstrated to reduce PHGDH expression **(Fig 5A, 5B)**. Further analysis of the combination therapy in 3D spheroids derived from a patient with BRAF^V600E^-driven ATC demonstrated similar anti-proliferative effect compared to single agent (**Fig 5D).**

**Figure 5.**
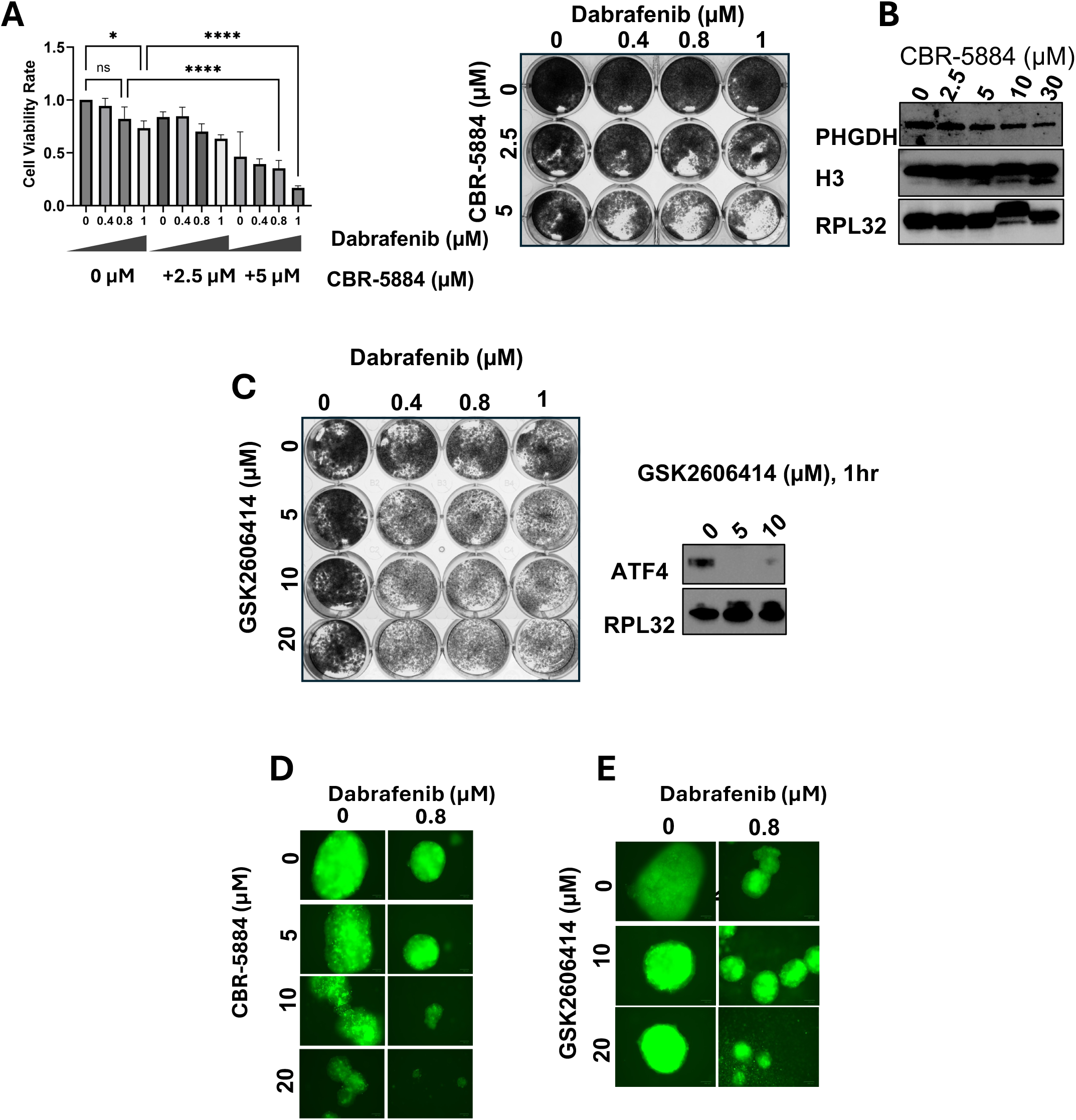
PHGDH and PERK inhibition mimics loss of TAZ in sensitizing ATC cells to dabrafenib. **A.** CBR-5884 sensitizes 8505c to dabrafenib and **B**. reduces PHGDH expression. **C**. GSK2606414 reduces ATF4 expression after 1 hour of treatment and sensitizes 8505c to dabrafenib. **D, E**. CBR-5884 and GSK2606414 sensitizes patient-derived spheroids (ATC-01) to dabrafenib after 10 days of treatment. *p<0.05, **p<0.01, ***p<0.001, ****p<0.0001.

To determine the role of PERK activation in response to ER stress and UPR and ATF4 activation upon dabrafenib treatment, PERK inhibitor GSK 2606414 was used. Although GSK414 impaired ATF4 expression, it did not affect tumor cells proliferation when used as a single agent. It did sensitize ATC cells to dabrafenib (**Fig 5C**). Similar findings were obtained in spheroids derived from an undifferentiated thyroid cancer patient sample (**Fig 5E**).

## Discussion

Anaplastic thyroid carcinoma (ATC) is a highly aggressive subtype of thyroid cancer, representing approximately 2% of cases, with a median survival of only 3–7 months. Conventional therapies offer limited efficacy, and patients with ATC often face a poor prognosis due to rapid tumor progression and distant metastases. Currently, no effective treatments have been shown to significantly improve overall survival in ATC patients.

UPR signaling is initiated in response to the accumulation of misfolded proteins in the ER [34]. Under ER stress conditions, the protein chaperone HSPA5 binds preferentially to misfolded proteins, thus releasing its inhibitory function on IRE1α, PERK and ATF6 [34]. To resolve the accumulation of misfolded proteins and restore proteostasis, activated PERK induces a transient inhibition of protein synthesis and a selective translation of ATF4 [35]. ATF4 translocated to the nucleus activates the transcriptional expression of essential genes for ER stress adaptation. These genes are involved in amino acid synthesis and transport, protein synthesis and folding, redox homeostasis, and cell death pathways [36–38].

ATF4-induced amino acid synthesis and transporter upregulation has been shown to be involved in rescuing the inhibition of protein synthesis during a prolonged ER stress by promoting mTOR signaling [39]. This increase in amino acid pool is an adaptive mechanism in response to ER stress and protein depletion.

Our findings show that dabrafenib triggers ATF4 expression and blocks protein synthesis. Consistent with these findings, previous studies performed in BRAFV600E-driven melanoma cells showed that BRAF inhibitor vemurafenib triggers ER stress and increases ATF4 expression by increasing the cytosolic Ca^2+^ [40]. Transcriptomic and proteomic analyses revealed increased of serine synthesis pathway (SSP) and serine transporters associated with ATF4 expression in response to the UPR. We also found that the rate limiting enzyme PHGDH is increased in dabrafenib-treated ATC cells in response to the ER stress and blockade of protein synthesis. In addition, the targeting of UPR sensors or downstream proteins sensitize ATC cells to BRAF inhibitor.

Increase in SSP enzymes, such as PHGDH, as demonstrated in our model, is the rate-limiting enzyme in serine de novo synthesis from the glycolytic pathway. However, several studies have shown that the SSP is not a limiting factor in all tumors [24, 41]. Our findings indicate that pharmacological inhibition of PHGDH at low concentrations exhibited a modest effect on cell growth but significantly sensitized ATC cells to dabrafenib.

Increased serine synthesis or uptake of exogenous serine from the tumor environment by solute carrier family 1 member 4 (SLC1A4) and solute carrier family 7 member 5 (SLC7A5) is necessary for therapy resistance in many cancers [42]. Non-small cell lung cancer (NSCLC) and melanoma cancer models treated with vemurafenib activated serine/glycine synthesis as a mechanism of acquired resistance to targeted therapy, and the silencing of PHGDH reversed resistance to therapy [43]. A recent study has shown a similar pattern in NRAS-mutated melanoma resistant to MEK inhibitor [44] .PHGDH plays a key role in ferroptosis by binding to PCBP2 and inhibiting its degradation, which in turn stabilizes SLC7A11 mRNA and makes cells resistant to ferroptosis [45]. Serine metabolism is a key factor in the maintenance of the redox homeostasis. Serine is a precursor of glycine and cysteine which are required for glutathione synthesis. Glutathione depletion inactivates glutathione peroxidase 4 (GPX4), a critical cellular antioxidant enzyme, triggering ferroptosis [46]. Taken together, our data suggest ATC cells exposed to dabrafenib hijack the UPR and subsequent SSP activation to survive BRAFV600E inhibitor treatment, by inactivating cell death pathways.

While most of the recent studies to improve response to BRAF inhibitors in ATC focused on drug repurposing, to our knowledge our study represents the first to utilize a novel strategy to identify genetic determinants of resistance to BRAFV600E inhibitor treatment in ATC models. Combining a CRISPR/KO screen with a CRISPR activation screen identified WWTR1, a gene coding for TAZ, as a key driver of resistance to BRAF inhibitor in ATC *in vitro* and in mouse model. Our screening results parallel previous studies demonstrating that dysregulated Hippo signaling pathway and subsequent activation of YAP/TAZ is a major resistance mechanism to targeted therapies, including EGFR inhibitor and MAPK inhibitors [47].

Functionally we demonstrate the role of TAZ in dabrafenib resistance using genetic models and pharmacological inhibitors for TAZ-TEAD binding. Mechanistically, we demonstrated that TAZ regulates ATC cell growth and modulates UPR signaling and protein synthesis. Furthermore, in the presence of dabrafenib, TAZ-depleted cells harbored reduced UPR signaling, partially restored protein synthesis, and reduced tumor growth *in vivo*. Our findings demonstrate that TAZ-mediated UPR confers resistance to BRAFV600E inhibitors (**Fig 6**). This conclusion is supported by various studies demonstrating the correlation between the UPR and resistance to chemotherapy in solid tumors. For instance, expression of UPR sensors and their downstream targets has been associated with shorter survival and higher risk of recurrence in patients with breast and colorectal cancers [48–50]. In many cancer types, drug-induced UPR leads to autophagy and cell death resistance and modulating UPR with pharmacological inhibitors sensitizes cancer cells to therapeutic agents such as gemcitabine or imatinib. Resistance of melanoma cells to the BRAFV600E inhibitor, PLX4032, is reversed by inhibiting the protein chaperone HSPA5 [51]. Taken together, our findings support the role of TAZ in resistance to dabrafenib. Blocking TAZ in combination with BRAFV600E inhibitor inhibit UPR signaling and its downstream effectors, thus leading to increased vulnerability to stress and cell death by ferroptosis. While the role of TAZ in modulating ferroptosis is molecular context- and cell type-dependent, a recent study on hepatocellular carcinoma (HCC) reported its implication in preventing ferroptosis in sorafenib-resistant HCC cells through its binding to ATF4 [52]. Several key proteins interacting with ATF4 such as NRF2, GPX4, SLC7A11, PDIA4 have been suggested as mediators of the role ATF4 plays in preventing ferroptosis in various cancer models [53]. However, in ATC, the precise mechanism leading to ferroptosis remains unelucidated, and warrants further investigations.

**Figure 6.**
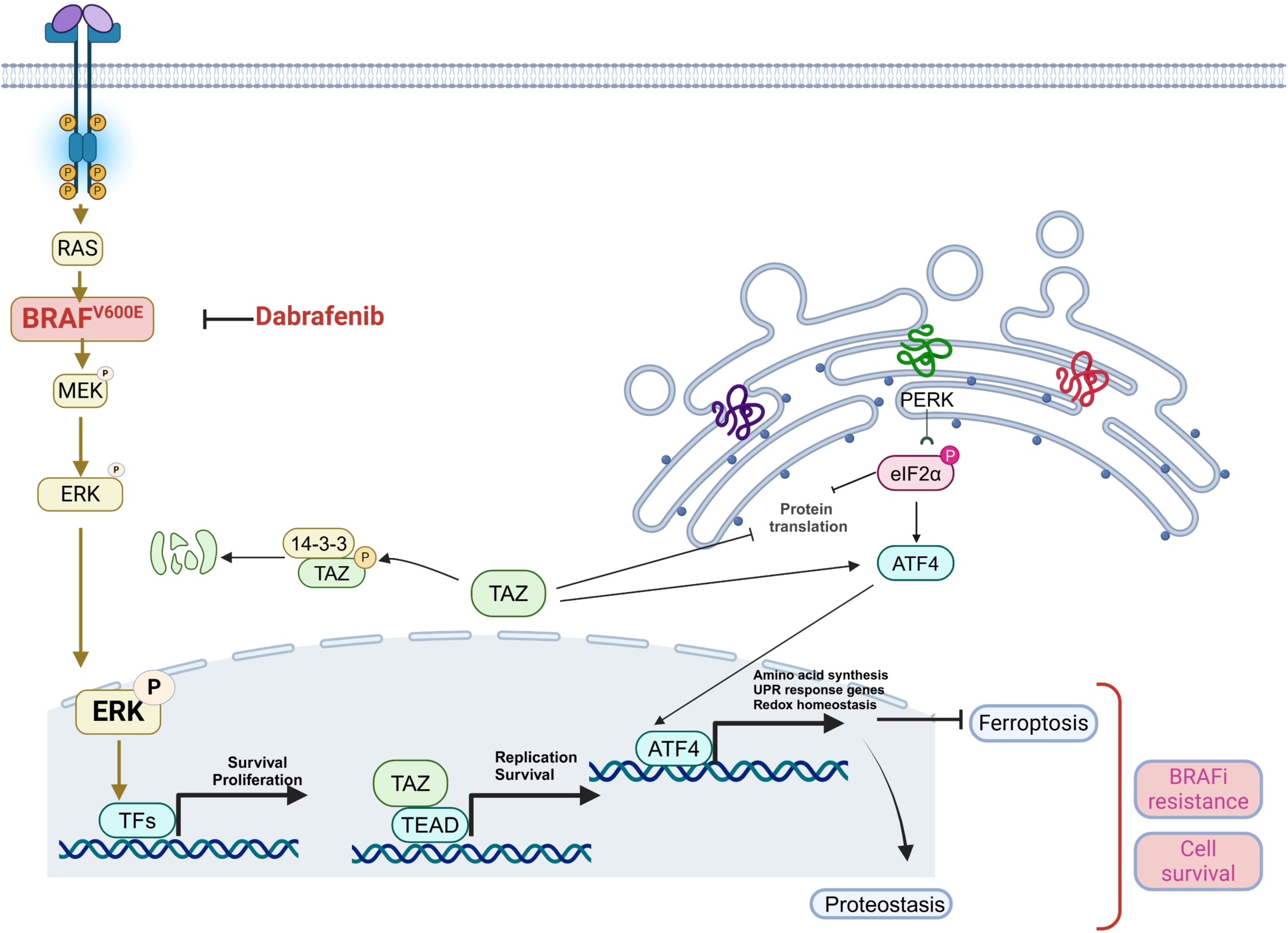
Working model describing the role of TAZ in dabrafenib resistant ATC. Image generated by Biorender.

Our study is the first to leverage CRISPR based screening to identify TAZ and the UPR as the genetic determinants of resistance to dabrafenib in undifferentiated thyroid cancer. Our findings corroborate the importance of TAZ and ER stress in resistance to targeted therapy. In addition, our results support the role for the SSP in cancer drug resistance as previously shown with other inhibitors such as sorafenib or sunitinib [54] [55]. The targeting of TAZ and BRAF^V600E^ is an attractive therapeutic strategy for ATC patients.

## Materials and methods

### Reagents

Dabrafenib (Cat. No. S2807), PHGDH inhibitor CBR-5884 (Cat. No. S9645), PERK inhibitor GSK2606414 (Cat. No. S7307), RSL3 (Cat. No. S8155) were purchased from Selleckchem. Pan-TEAD inhibitor GNE7883 (Cat. No. HY-147214) was purchased from MedChem Express. Thapsigargin was purchased from Selleckchem.

### Cell Culture

The human anaplastic thyroid cancer cell line 8505c (Sigmal Millipore) was grown at 37°C with 5% CO_2_ in Dulbecco’s Modified Eagle Medium (DMEM) (HyClone) supplemented with 10% Fetal Bovine Serum (Gemini Bio) and 1% Penicillin Streptomycin (Gibco). The patient-derived anaplastic thyroid cancer cell lines ATC01 was grown at 37°C with 5% CO_2_ in Dulbecco’s Modified Eagle Medium/Nutrient Mixture F-12 (DMEM/F-12) (HyClone) supplemented with 15% Fetal Bovine Serum and 1% Penicillin Streptomycin as described by Hu et *al* [15].

### RNA Sequencing

Sequencing data were processed using CCBR RENEE Rnaseq pipeline (https://github.com/CCBR/RENEE) using hg38 genome (Gencode Release 41). This work utilized the computational resources of the NIH HPC Biowulf cluster (http://hpc.nih.gov).

Downstream analysis and visualization were performed within the NIH Integrated Analysis Platform (NIDAP) using R programs developed by a team of NCI bioinformaticians on the Foundry platform (Palantir Technologies), the code will be public. After RNA-Seq FASTQ files were aligned to the reference genome using STAR [16] and raw counts data produced using RSEM [17], the gene counts matrix was imported into the NIDAP platform, where genes were filtered for low counts (<1 cpm) and normalized by quantile normalization using the limma package [18]. Differentially expressed genes were calculated using limma-Voom [19]. GSEA was performed using fgsea package (6) and MSigDB v6.2.

### In vivo Studies

Animal studies were approved by the National Cancer Institute’s Institutional Animal Care and Use Committee. The mice were maintained according to the guidelines of the National Cancer Institute’s Animal Research Advisory Committee. Briefly, 1.6 million 8505C cells depleted for TAZ or control 8505c cells were injected subcutaneously into the flank of Cg-*Prkdc^scid^ Il2rg^tm1Wjl^*/SzJ mice. Mice were randomized and treated daily by gavage with either 15mg/kg dabrafenib or vehicle. Tumor size was measured once a week for the duration of the experiment. For survival analysis, animals were euthanized once they showed at least one of the humane euthanasia criteria.

### Statistical Analyses

Statistical analyses were performed using GraphPad Prism 10 software (GraphPad). A p-value less than 0.05 was considered statistically significant. Data are presented as mean ± SD.

## Conflict of interest

the authors declare no conflict of interest

## Supporting information

This file contains all supplementary materials and data

## Acknowledgements

This research was supported in part by the Intramural Research Program of the National Institutes of Health (NIH), National Cancer Institute, and the Center for Cancer Research. This work used the computational resources of the NIH HPC Biowulf cluster (https://hpc.nih.gov). We are grateful to our colleague Ms. Elena Kuznetsova who passed away recently.

## Data Availability Section

All data and codes will be made available in a link

## Notes

### Competing Interest Statement

The authors have declared no competing interest.

