## Supplementary material for "CRISPR-Based Gene Dependency Screens reveal Mechanism of BRAF Inhibitor Resistance in Anaplastic Thyroid Cancer": This file contains all supplementary materials and data

**Supplementary materials and methods**

**CRISPR/Cas9 Knockout and Activation Library Screens**

In this study, a focused CRISPR gene library was used to identify genes responsible for BRAF inhibition resistance in the human anaplastic thyroid cancer cell line 8505c.

*Library Design*

The library was designed by first generating a list of genes in BRAFV600E-mutated cell lines with a negative CHRONOS gene effect score in Cancer Dependency map (DepMap). This list was then combined with a list of differentially expressed genes in dabrafenib treated cells, generated by performing RNA sequencing on 8505c cells treated with dabrafenib for 14 days and data were analyzed using Partek Flow software. After combining these gene lists and removing noncoding RNAs (ncRNAs) and pseudogenes, we created a focused CRISPR gene library of 2,050 genes used for the knockout and activation screens.

*Construction of CRISPR-KO/CRISPR-activation libraries*

For the generation of the targeted CRISPR KO library, guide RNAs were obtained from multiple sources. Namely, from the Brunello genome wide library as well as designed using a combination of the GuideScan algorithm and sgRNA Scorer [1-3]. For the CRISPR activation library, guide RNAs were pulled from the Calabrese A and B libraries (PMID: 30575746) as well as the hCRISPRa-V2 library (PMID: 27661255). In total, up to 6 guides per gene were designed for both libraries.

Guide RNA sequences, along with appropriate flanking sequence to facilitate cloning, were synthesized as single stranded DNA (ssDNA) oligo pools from Twist Biosciences. For the CRISPRko library, the lenti-guide-puro (Addgene-52963) [4] was used as the backbone and for the CRISPRa library, the lenti sgRNA(MS2)_puro (Addgene-73797) [5] backbone was used. lenti sgRNA(MS2)_puro optimized backbone (Addgene plasmid # 73797 ; http://n2t.net/addgene:73797 ; RRID:Addgene_73797) and lentiGuide-Puro (Addgene plasmid # 52963 ; http://n2t.net/addgene:52963 ; RRID:Addgene_52963) were gifts from Feng Zhang. Oligo pools were then amplified minimally to generate double stranded PCR product. Six 20 uL HiFi Builder reactions with 100 ng of BsmBI digested backbone and 10 ng of dsDNA PCR product per reaction were then set up for each library. Assembly products were then pooled, column purified and eluted in 13 uL of elution buffer. Eluted product was the electroporated into Lucigen Endura cells and plated over 10 – 15 cm agar plates and grown at 30C. Colonies were then scraped, pelleted, and then maxiprepped using the Genejet Endo-free maxiprep kit. Libraries were then sequenced for representation using the Illumina NextSeq.

*CRISPR Screens*

Cas9-expressing cells for the KO screen and dCas9-synergistic activation mediator (SAM) complex expressing cells for the activation screen were established by lentiviral transduction using the following plasmids: Addgene #52962, #61425, #89308.

8505c-Cas9 and 8505c-dCas9-MPH+ cells were transduced with the focused CRISPR library, which contains 12,344 gRNAs targeting 2,050 human genes (~6 gRNAs per gene and 150 control gRNAs targeting intergenic region), at a low MOI (~0.4) to ensure effective barcoding of individual cells. The transduced cells with CRISPR KO-or CRISPR-activation were selected with 0.5 µg/mL of puromycin for 7 days to generate a mutant cell pool. The mutant cell pool was divided into 3 groups; a subset of cells was frozen to serve as genomic DNA control for data analysis, while the rest were treated with either vehicle or dabrafenib (0.4 µM) for 14 days. After treatment, at least 6.2 million cells were collected for genomic DNA extraction to ensure over 500X coverage of the CRISPR gene library.

*Method for sgRNA sequencing and data analysis*

PCR primers were designed with IDT PrimerQuest. The PCR amplification was done with Takara Titanium® Taq DNA Polymerase (Takara, Cat#639242) according to the manufacture's protocol. PCR products for the same sample were pooled then purified using SPRI beads. Qubit HS DNA kit was used to quantitate the library concentration. The library molar concentration is determined with Qubit concentration and actual PCR product length. The libraries were sequenced on Illumina NextSeq2000 with 100 +8 cycles. The sequence reads were analyzed with Perl script to known and unknown sgRNA inserts. The sgRNA sequences were amplified using NEBNext®High-Fidelity 2X PCR Master Mix and subjected to Next Generation Sequencing. The sgRNA read count and hits calling were analyzed using the *MAGeCKFlute* pipeline (version mageck-0.5.9.4).

**Gene Essentiality Analysis and Public Datasets**

For essentially score, normalized growth reduction resulting from gene inactivation of WWTR1 from DepMap (Public 22Q4 version) and BRAF somatic mutation information from cBioPortal we used. P-values were calculated using a one-sided *Wilcoxon rank-sum* test.

GSE33630 from Gene Expression Omnibus was used to analyze WWTR1 and ATF4 expression in normal thyroid tissue, papillary thyroid cancer and anaplastic thyroid cancer cohort. Gene expression profile data was downloaded from the Gene Expression Omnibus database (https://www.ncbi.nlm.nih.gov/geo/query/acc.cgi?acc=gse33630) from the National Center for Biotechnology Information based on the platform of the GPL570 Affymetrix Human Genome U133 Plus 2.0 array.

**Generation of Stable Knockdown and Knock-out Cell Lines**

TAZ knockdown cells were generated using *TAZ*-targeting shRNAs (MilliporeSigma TRCN0000019469 and SHC001 control plasmid) 8505c were infected with viral supernatant containing the shTAZ or shControl particles. 72hrs post infection selection antibiotic was added. Newly generated cell lines were selected for 7 days with Puromycin 0.5 µg/ml. The cells were then harvested and confirmed for depletion by Western blot. For CRISPR/cas9 KO cells, 8505c-Cas9+ cells were infected as described above with viral supernatant containing the pKLV2-U6gRNA5(hWWTR1-W1K)-PGKpuro2AmCherry-W (Addgene 163176),pKLV2-U6gRNA5(hWWTR1-W2B)-PGKpuro2AmCherry-W( addgene163177). pKLV2-U6gRNA5(BbsI)-PGKpuro2AmCherry-W (Addgene 67977) was used a control.

**TAZ Overexpression**

8505c were infected with viral supernatant containing an optimized version of TAZ. Infected cells were selected with 300 µg/mL of Geneticin Selective Antibiotic (Gibco, Cat No 10131035).

**siRNA Transfection**

8505c cells were reverse transfected with an siRNA duplex using Lipofectamine RNAiMAX Reagent (Invitrogen, Cat No 13778075) according to the manufacturer's instructions for 48 hours. siATF4 and siPHGDH were obtained from Thermo Fisher Scientific.

**RNA extraction**

RNAs were extracted using RNeasy kit (Qiagen) according to the manufacture's protocol. RNA concentrations were determined using a nanodrop and RNA quality was assed using a bioanalyzer. Samples were used for RNA sequencing or RT-PCR. For RNA seq we used the NextSeq 2000 XLEAP P4 200 cycle kit,

**RT-PCR**

The mRNA expression of WWTR1, ATF4, PHGDH, PSAT1, PSPH were determined by RT-PCR. RPLPO was used as a housekeeping gene. RNA was extracted using a RNeasy Mini Kit (Qiagen). RNA yield was determined using a NanoDrop One spectrophotometer (Thermo Scientific). cDNA was synthesized from 500ng of total RNA using a High-Capacity cDNA Reverse Transcription Kit (Thermo Fisher Scientific). PCR was performed using TaqMan Gene Expression Master Mix according to the manufacturer's protocol.

**Western Blot**

The protein expression of TAZ, ATF4, PHGDH, phospho-MEK, and MEK was determined by Western blotting. RPL32, H3, and vinculin were used as loading controls. The protein lysate was extracted using a lysis buffer containing 50mM Tris-HCl, 150mM NaCl, 1% Triton X-100, supplemented with protease and phosphatase inhibitors. The total protein concentration was determined using the Pierce BCA Protein Assay Kit (Thermo Scientific, Cat. No. 23227). 15-30µg of total protein lysate was loaded into a 4-15% Criterion TGX Stain-Free Precast Gel (Bio-Rad) and then transferred onto a PVDF membrane. Membranes were immunostained with one of the following primary antibodies (1:1000 in 2.5% BSA) overnight at 4ºC: TAZ (Cell Signaling Technology 72804S), ATF4 (Cell Signaling Technology 11815S), PHGDH (Cell Signaling Technology 66350S), phospho-MEK1/2 (Cell Signaling Technology 9121S), MEK1/2 (Signaling Technology 9102S), RPL32 (Novus biologicals, NBP3-25485), and H3 (Cell Signaling Technology 4499S).

For protein synthesis assay, cells were incubated with puromycin at 10µg/ml for 10 minutes. Total protein lysates were then analyzed by western blot and probed with anti-puromycin antibody (MilliporeSigma, Cat. No. MABE343).

**Flow Cytometry Assays**

Cells were seeded in T25 flasks, after 24 hours, they were incubated with dabrafenib for 7 days. After treatment, cells were incubated in 10µM BODIPY 581/591 C11 Lipid Peroxidation Sensor (Invitrogen, Cat No D3861) for 30 minutes at 37ºC, or cells were incubated in 5µM CellROX Green Reagent (Invitrogen, Cat No C10444) for 30 minutes at 37ºC. Samples were trypsinized, centrifuged, and resuspended in 1mL PBS. For the BODIPY assay, fluorescence was detected at 581/591 nm (Ex/Em) for the reduced dye and at 488/510 nm (Ex/Em) for the oxidized dye using a BD LSRFortessa flow cytometry analyzer. For the CellROX Green assay, fluorescence was detected at 485/520 (Ex/Em) using a BD LSRFortessa. RSL3 and $H_2O_2$ were used as positive controls, respectively.

**Clonogenic Assays**

Cells were seeded into 12-well plates and treated for seven days, refreshing the vehicle or drug containing medium every other day. Cells were fixed in 100% methanol for 15 minutes and stained with a solution of 0.4% (w/v) Brilliant Blue (B0770-25G) in $H_2O$ for 45 minutes. Colony formation was quantified using the Image Lab software (Bio-Rad).

**Immunoprecipitation**

Treated cells were lysed using a lysis buffer containing 25mM Tris-HCl pH 7.4, 150mM NaCl, 1% Triton X-100, and 5% glycerol. 750μg of protein were incubated in the appropriate primary antibody overnight at 4ºC. Protein A/G magnetic beads were incubated and rocked in lysis buffer overnight at 4ºC. The beads were then washed twice and resuspended with PBS. 20μL of A/G beads were incubated and rocked with the protein-antibody solution for 1 hour at 4ºC. The samples were washed six times with the lysis buffer and then resuspended in 2x Laemmli sample buffer with DTT. After heating the solution at 99ºC, the sample was isolated from the beads and loaded into a gel for analysis alongside a total protein lysate input. Immunoblot for IP samples were incubated with Veriblot-HRP secondary antibody (abcam, Cat. No. ab131366).

**Cell Viability and Apoptosis Assays**

Cells were seeded at 1,500 cells/well into a 96-well plate and treated for seven days, refreshing the vehicle or drug containing medium every other day. Cell viability was determined by the CellTiter-Glo Luminescent Cell Viability Assay (Promega, Cat. No. G7571) according to the manufacturer’s instructions. Caspase dependent apoptosis was determined using Caspase 3/7 Glo assay (Promega, Cat. No. G8090).

For cell proliferation assay of ATC01 patient-derived spheroids, cells were seeded at 4000 cells/ well of a 96 wells low attachment plate, once spheroids are formed, drugs were added and renewed every 48hrs, cell titer glo was used to measure cell growth.

The Calcein-AM assay (Calcein AM, 354216, Corning) was used to assess the toxicity of the drugs combination in ATC-01 spheroids. 30,000 cells were seeded in each well of low-attachment 24 wells plates. Once spheroids are formed, drugs were added and renewed every 48hrs for the duration of the experiment. At the end of the experiment, we aspirated the cell culture medium and incubated the cells with PBS containing 1uM of Calcein AM, for 1h at 37 °C in 5% $CO_2$. Green fluorescence was then detected using a fluorescence cell imager (ZOE Fluorescent cell imager, Biorad).

#### Proteomics

*Digestion and Sample Processing*

Cell pellets were resuspended in 200 uL EasyPep™ (Thermo Scientific) Buffer with 1 uL nuclease and Phos Stop. BCA was used to estimate protein concentration and 100 ug of each sample was combined with 50 uL Reduction and 50 uL alkylation solution. 6 ug of Trypsin/LysC was added to each sample and allowed to digest overnight at 37°C. The next day 200 ug TMT label was added to each sample and allowed to label for 1h at 25°C. Reactions were quenched with 50 uL of 5% hydroxylamine, 20% Formic acid for 10 minutes and samples were combined and cleaned up using the EasyPep™ (Thermo Scientific) Maxi Kit. Eluted samples were dried in the speed-vac

*LC/MS Analysis*

The sample was resuspended in 50 uL of 0.1% FA and 5 uL was loaded onto a Dionex U3000 RSLC attached to an Orbitrap Eclipse (Thermo) equipped with a FAIMS and EasySpray ion source. Solvent A was comprised of 0.1% FA and Solvent B was 0.1% FA in 80% CAN. The gradient pump was run at 300 uL/min with an LC gradient of 5-7%B for 1 min, 7-30%B for 83 min, 30-59%B for 25 min, 50-95%B for 4min, 95%B for 7 min then a re-equilibration of the column at 5%B for 17 min. The MS was run in the TopSpeed method with four FAIMS compensation voltages (45, 55, 65, 75 or 50, 60, 70, 80). Spray voltage was set at 2200V and the ion transfer tube was at 300°C. MS1 scans were acquired in the Orbitrap at 120,000 resolution, AGC of 4e5, and max injection time of 50ms in a mass range of 375-1600 m/z. MS2 scans were acquired in the Orbitrap using the TurboTMT method with a resolution of 15,000, intensity threshold 2.5e4, AGC 5e4, HCD energy 30%, isolation window 1.6 m/z and charges of 2-5 for MS2 selection. Monoisptopic Precursor selection (MIPs), Easy-IC for internal calibration and advanced peak determination were enabled.

*Database search and post-process analysis*

The MS files were searched in Proteome discoverer using the Sequest node. Data was searched against the Uniprot human data base using full trypsin digestion with a max 2 missed cleavages, Min peptide length 6, MS1 mass tolerance of 10ppm, MS2 tolerance of 0.02 Da, variable modification of oxidation on methionine and static modifications of carbamidomethyl on cysteine, and TMTpro on lysine and peptide N-terminus. Percolator was used for FDR analysis and TMTPro reporter ions were quantified using the Reporter Ion Quantifier node and normalized on total peptide intensity on each channel.

#### Supplementary Figure 1

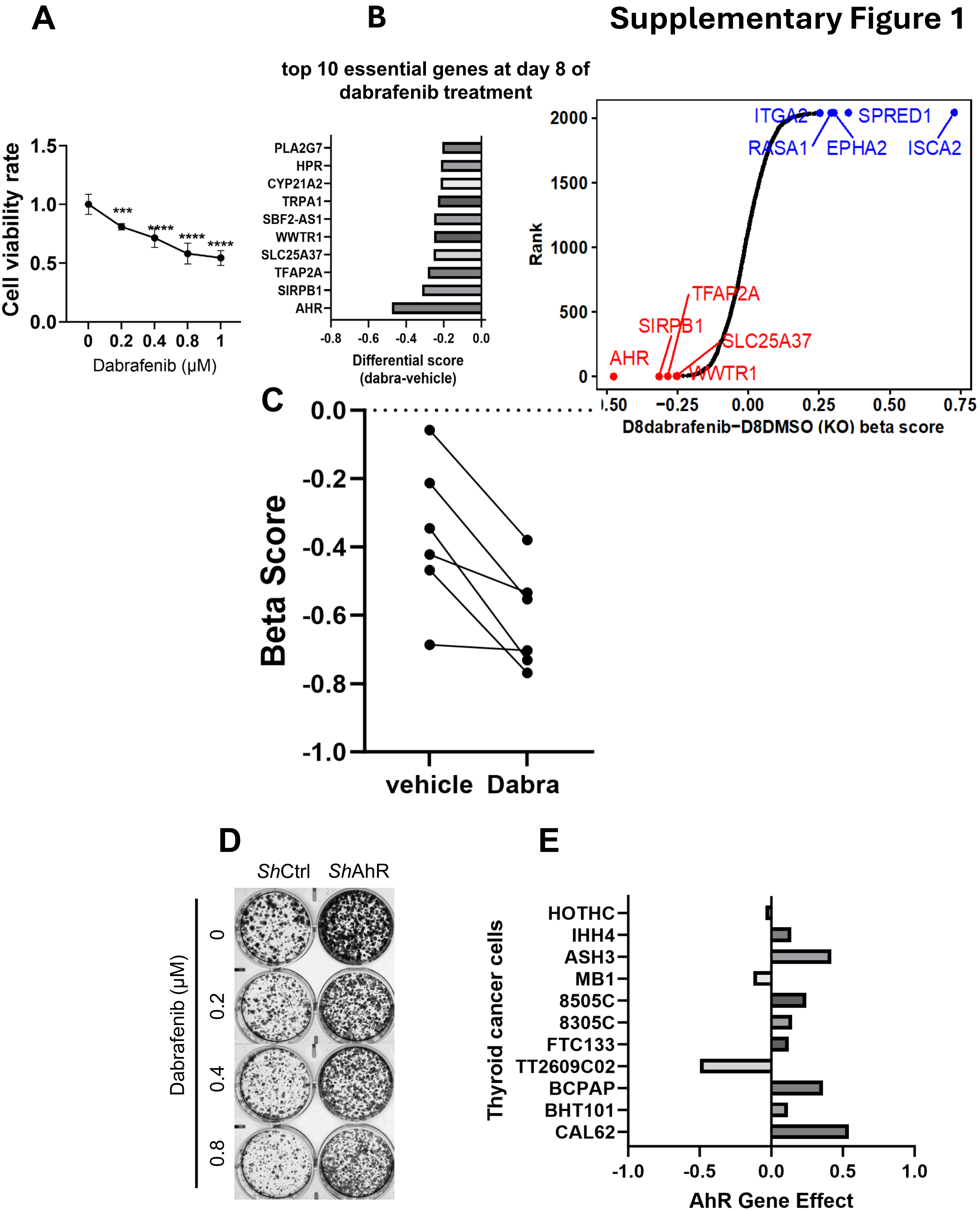

**Supplementary Figure 1: A.** cell growth analysis in response to increasing doses of dabrafenib at 7 days. **B**. Top ten essential genes from CRISPR/Cas9 KO screen at days 8 of KO screen. **C**. sgRNA targeting WWTR1 were depleted in dabrafenib treated cell at day 8. **D**. Top hit of both CRISPR/KO screen AhR, increased cell proliferation. **E**. DepMap analysis showed a positive gene effect with AhR KO in most of thyroid cancer cells. *$p<0.05$, **$p<0.01$, ***$p<0.001$, ****$p<0.0001$

### Supplementary Figure 2

A

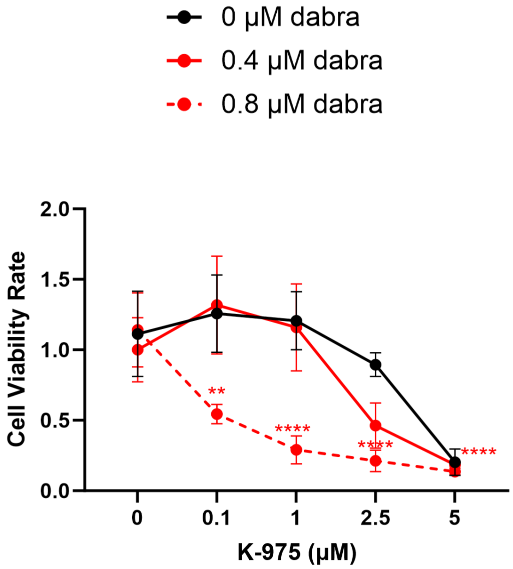


B

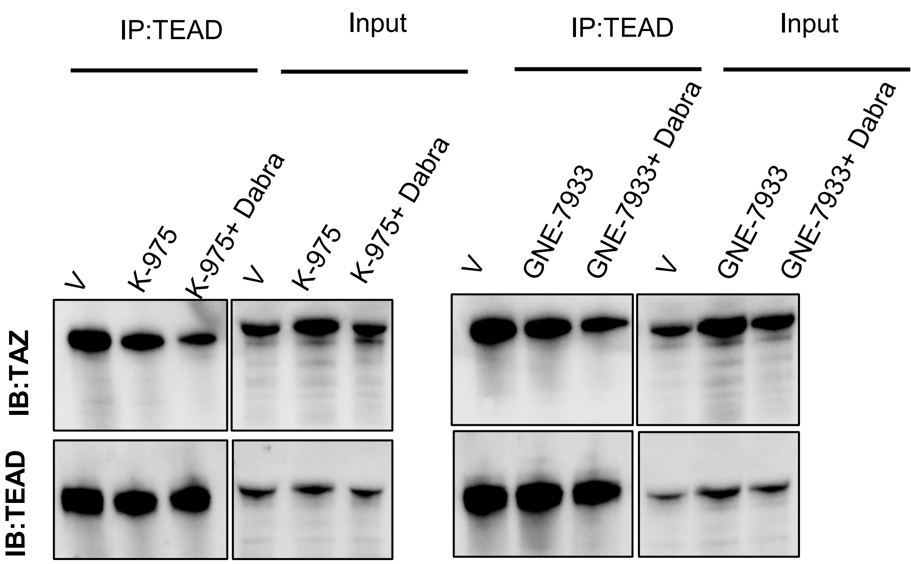

**Supplementary Figure 2: A**. K-975 sensitized ATC cells to dabrafenib. **B.** Combination of dabrafenib with K-975 or GNE-7933 reduced the interaction of TAZ with Pan-TEAD compared to K-975 or GNE-7933 alone

### Supplementary Figure 3

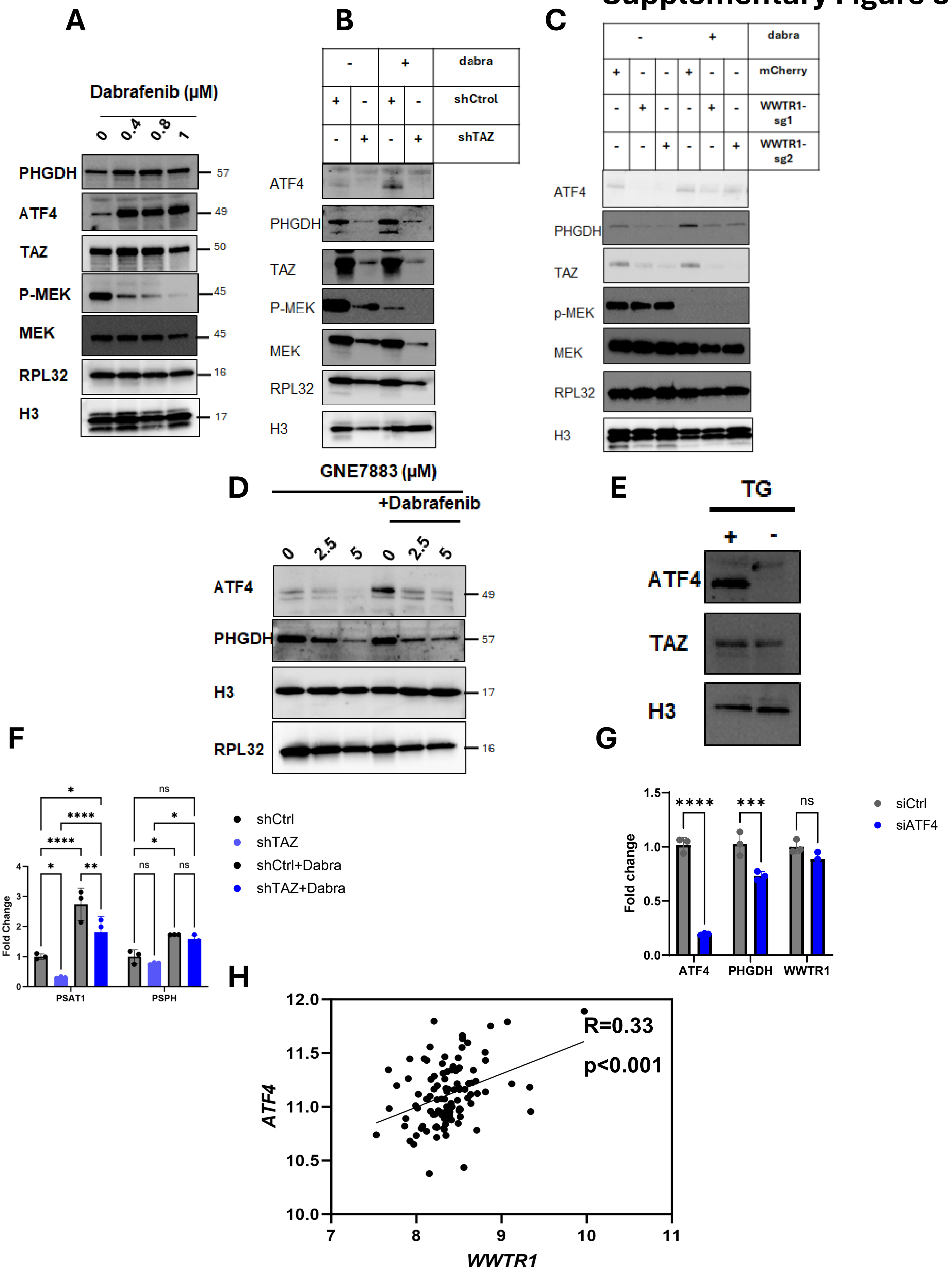

**Supplementary Figure 3**: **A**. UPR protein levels in response to increasing concentrations of dabrafenib. 7 days of dabrafenib treatment increases ATF4 and PHGDH and has a mild effect on TAZ expression. Dabrafenib reduces p-MEK in a dose dependent manner. **B**, **C**. Cells depleted for TAZ (shTAZ and TAZ KO) were treated with dabrafenib for 72hrs. ATF4, PHGDH and p-MEK were analyzed. TAZ depletion was confirmed in shTAZ and TAZ KO cells**. D.** The effect of GNE-7833 mimics that of TAZ depletion on dabrafenib-induced ATF4 and PHGDH. **E.** TAZ and ATF4 in Thapsigargin (TG) treated cells for 18hrs**. F**. RT-PCR analysis of PSAT1 and PSPH in dabrafenib treated cells with and without TAZ depletion **G.** RT-PCR analysis of ATF4, PHGDH and TAZ in siCtrl and siATF4 cells. **H.** Spearman correlation of *ATF4* and *WWTR1* in all samples of GSE33630.

### Supplementary Figure 4

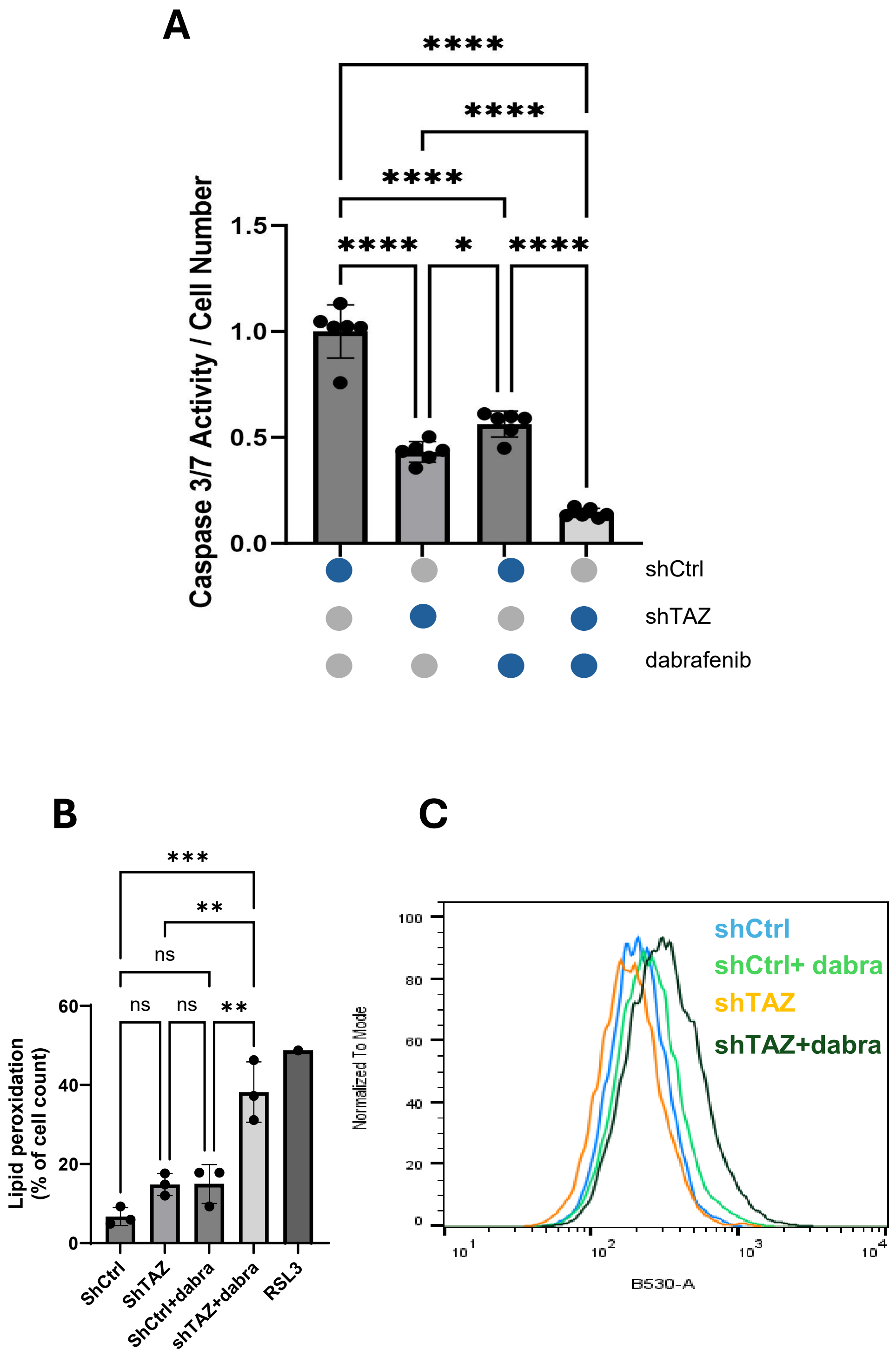

**Supplementary Figure 4. A.** Caspase3/7 activity in shCtrl and shTAZ cells treated with vehicle or dabrafenib at 7 days. **B.** Lipid ROS level was analyzed using BODIPY in shCtrl, shTAZ, shCtrl+dabrafenib and shTAZ+dabrafenib treated cells**.** RSL3 was used as a positive control. Lipid peroxidation for all replicates is summarized in the histogram. **C**. ROS level was measured using CellROX Green in the same conditions as **B**. *p<0.05, **p<0.01, ***p<0.001, ****p<0.0001
